# Selection of popcorn genotypes tolerant to *Spodoptera frugiperda* and key traits related to the identification of tolerance

**DOI:** 10.1101/2021.01.04.425203

**Authors:** Amanda Tami Kuroda, Jocimar Costa Rosa, Luiz Felipe Antunes de Almeida, Giovana Dal Lago Garcia, Gustavo Arana Demitto, Renata Maria Bento de Souza, Fernando Alves de Albuquerque, Carlos Alberto Scapim

## Abstract

The *Spodoptera frugiperda*, is one of the most deleterious pests of popcorn and the identification of tolerant genotypes is determinant in breeding programs. The objective of this study was to select popcorn genotypes tolerant to *S. frugiperda* and the key traits related to the identification of tolerance. The popcorn varieties UEM J1, Composto Márcia, Arachida, Composto Gaúcho and Zapalote Chico (resistant check) were evaluated in a completely randomized design with 100 replications. The experimental unit consisted of one Petri dish, containing plant material and a larva. The following traits were evaluated: larval stage duration (LSt), food intake weight (IW), final larva weight (FW), mean larva weight (MW), feces (F), assimilated (A) and metabolized food weight (M), relative consumption rate (RCR), relative metabolic rate (RMR), relative growth rate (RGR), conversion efficiency of ingested food (CEI), apparent digestibility (AD), conversion efficiency of digested food (CED) and leaf area consumed (LAC). The diagnosis of multicollinearity, analysis of canonical variables, genetic divergence, hierarchical clustering, factor analysis and canonical correspondence analysis were carried out to perform multivariate analysis. After the multicollinearity test, the traits FW, IW, RCR, AD and LAC were maintained for further analysis. Variety Arachida was considered tolerant to *S. frugiperda* and can be used in the future as a source of favorable alleles to breed tolerant popcorn hybrids. The traits relative consumption rate, apparent digestibility and leaf area consumed were considered key traits in the identification of tolerance against *S. frugiperda* in popcorn genotypes.

## Introduction

The fall armyworm, *Spodoptera frugiperda*, is considered one of the most deleterious pests of maize in all maize-producing regions [1]. In the tropics, it causes significant economic damage because it feeds on maize from the seedling to the reproductive stage [2]. In popcorn, which has been less the focus of breeding efforts than field corn, the damage tends to be more severe, since no tolerant genotypes are available on the market. Considering the increasing nationwide and worldwide demand for and importance of popcorn [3], the possibility of control by the development of tolerant genotypes must be investigated and improved, since the number of insecticide applications has increased considerably over the years [4, 5], resulting in excessive costs and increased environmental risks.

In view of the concern to reduce or even eliminate insecticide applications and only a few of studies involving *S. frugiperda* tolerance in popcorn, breeding programs ought to intensify studies to the selection of tolerant popcorn genotypes, as well as to elucidate key traits related to the identification of tolerance [6, 7].

Although some traits related to identification of *S. frugiperda* tolerance, e.g., larval stage duration, final and mean larva weight, consumption rates and leaf area consumed have been studied in maize breeding programs [7], they have not yet been conclusively proven and used as key traits. A likely reason may be that the trials were analyzed based on a univariate and inconclusive approach. The use of multivariate techniques is required, which can be an adequate and more efficient tool in the analysis of tolerance-related data in popcorn [8-10].

Resistance is the set of physical, chemical and morphological traits that will negatively affect the insect’s oviposition and feeding behavior, while tolerance is the set of traits that will cause plants to withstand the attack of insects without substantial reductions in productivity in comparison with other susceptible genotypes [8, 9, 11].

In this study, the hypothesis was proposed that tolerance to *S. frugiperda* can be found in popcorn germplasm from tropical regions and that multivariate analysis can discriminate the main traits related to the identification of tolerance. The objective of this study was to select popcorn genotypes tolerant to *S. frugiperda* and the key traits related to the identification of tolerance.

## Material and methods

The trial was carried out in Maringá, Paraná, Brazil, to evaluate five popcorn varieties (Table 1) for tolerance to fall armyworm (*Spodoptera frugiperda)*, as well as the key traits related to the identification of this tolerance.

**Table 1.** Description of the genotypes selected for study.

| Code | Genotype | Origin | Grain type | Genetic base |
| --- | --- | --- | --- | --- |
| 1 | UEM J1 | LIV | Popcorn | Open-pollinated variety |
| 2 | Zapalote Chico | CIMMYT | Field corn | Open-pollinated variety |
| 3 | Composto Márcia | MABH | Popcorn | Open-pollinated variety |
| 4 | Arachida | LIV | Popcorn | Open-pollinated variety |
| 5 | Composto Gaúcho | MABH | Popcorn | Open-pollinated variety |

LIV: local landrace varieties, MABH: mixture of American and Brazilian hybrids.

The popcorn genotypes used in the trial consisted of plants of the varieties UEM J1, Composto Márcia, Arachida, Composto Gaúcho and Zapalote Chico, grown in a greenhouse. The first four evaluated genotypes were developed by the Specialty Corn Breeding program of the State University of Maringá. The variety used as check, Zapalote Chico, was introduced from Central América and has tolerance against fall armyworm on the basis of antixenosis (alimentary avoidance) and antibiosis (lower insect survival after feeding on host tissue) [12, 13].

The plants were grown in a greenhouse with automatic irrigation. Crop management was carried out in accordance with the recommendations for corn culture [16]. The seeds were separated and three sown in each pot, which contained soil and substrate (3:1). After sowing, the pots were irrigated daily and side-dressed with urea (45% N). The other cultural treatments were applied as required for full crop development, without using any other chemical product, so as not to affect larva growth.

The corn leaves used to feed the larvas were collected when the plants were in the eight-leaf (V8) stage, so that all genotypes were evaluated when the plants were in the same developmental stage [6, 7, 17].

The insects required to initiate the trial were hatched from *S. frugiperda* eggs donated by EMBRAPA Soybean, in Londrina, Paraná, and from eggs collected in corn fields on the Experimental Farm of Iguatemi - Maringá. The larvas hatched from these eggs were fed an artificial diet and three generations were reared for our use in the trial.

The laboratory trial was carried out in an air-conditioned chamber at 25°C±1, air humidity of 70% ± 10 and a 12h photoperiod. Each experimental unit consisted of a sterile acrylic Petri dish (diameter 9.0cm, height 1.5cm), lined with filter paper moistened with distilled water to maintain the leaf turgor, containing only one larva per dish, to avoid insect cannibalism. Each treatment consisted of three Petri dishes with moist filter paper and plant material, to calculate the water loss.

The trial was conducted in a completely randomized design, with five treatments and 100 replications.

After hatching of the fourth larva generation raised on artificial diet, they were distributed in separate Petri dishes. The filter paper was changed, plant material was supplied to feed the larvas and the biological parameters were evaluated daily. Plant material of the same genotype was continuously supplied until the end of the larval stage.

The traits were evaluated as proposed by [18] with changes made by [19] as follows:

> Larval stage duration: LSt
>
> Food intake weight: IW
>
> Final larva weight: FW
>
> Mean larva weight during T: MW
>
> Feces: F;
>
> Assimilated food: A = I – F
>
> Metabolized food: M = A - FW
>
> Relative consumption rate: RCR = I/(MW * T)
>
> Relative metabolic rate: RMR = M (MW * T)
>
> Relative growth rate: RGR = FW * (MW * T)
>
> Conversion efficiency of ingested food: CEI = (FW/I) * 100
>
> Apparent digestibility: AD = (H/A) * 100
>
> Conversion efficiency of digested food: CED = FW/A * 100
>
> Leaf area consumed: LAC = I/SDm, where SDm: Mean surface density.

Food consumption and use were evaluated daily throughout the larval stage, by weighing the fresh food weight, leftover food weight, feces weight and larva weight. The values were measured for each larva and the mean for each repetition was calculated according to the number of live larvas on the day of the evaluation. The final data were obtained from the mean of the 100 replications per treatment.

The larva weight was measured directly by individual weighing during the whole larval stage. The daily weighing of the larvas was only initiated after the 5^d^ after egg hatch. Food intake weight (IW) was calculated indirectly, by subtracting of the statistically corrected leftover food (Lc) weight from the weight of the supplied food (SF) on the day before.

The excreta were collected and weighed individually for each larva during the entire larval stage to obtain the total weight of feces produced (F). The leaf area consumed (LAC) was calculated indirectly from the relationship between food intake (I) and mean surface density (SDm).

The assumptions of normality of residuals and homogeneity of variances were evaluated by the Shapiro-Wilk and Levene tests, respectively. The data were processed statistically by multivariate analysis of variance (MANOVA), using the statistical software Genes [20] integrated with R software [21].

The Variance Inflation Factor (VIF) and the Condition Index (CI) were used as criteria to assess the degree of multicollinearity between the predictive traits. Variance inflation factors of > 10 are generally considered evidence of substantial multicollinearity and normally the reason for the removal of certain predictors. In addition, multicollinearity is considered weak when the CI is less than 100 [14, 16].

Genetic divergence between the genotypes was evaluated by UPGMA (Unweighted Pair-Group Method using Arithmetic Averages), based on Mahalanobis’ distance. The groups were established according to the methodology proposed by [22]. Then, canonical variable analysis was carried out with clustering by the Tocher method, factor analysis and later canonical correspondence analysis [23].

Factor analysis was performed considering all evaluated traits [24]. The factorial loads were extracted by the principal component method, and the factors established by varimax rotation. In this study, factor loads above 0.90 as well as the highest values for the community were considered [25]. Canonical correspondence analysis was performed as described by [26].

The analyses were performed using the statistical software Genes [20] and SAS [27] at 1% probability.

## Results and discussion

Normality (p > 0.01) and homogeneity of variances (p > 0.01) were reported for all evaluated traits. The MANOVA test showed significant differences between the mean vectors of the genotypes for all evaluated traits (p <0.01), indicating the existence of genetic variability.

Multicollinearity among traits was assessed by the criteria VIF and CI. Traits with VIF and CI values greater than 10 and 100, respectively, are generally considered evidence of substantial multicollinearity among variables and makes the removal of these predictors necessary [28, 15]. The traits LSt, MW, F, A, M, RMR, RGR, CEI and CED had high VIF and CI values and were therefore eliminated. On the other hand, the traits FW, IW, RCR, AD and LAC had VIF and CI had values below 10 and 100, respectively, and were therefore maintained for subsequent analyses (Table 2).

**Table 2.**
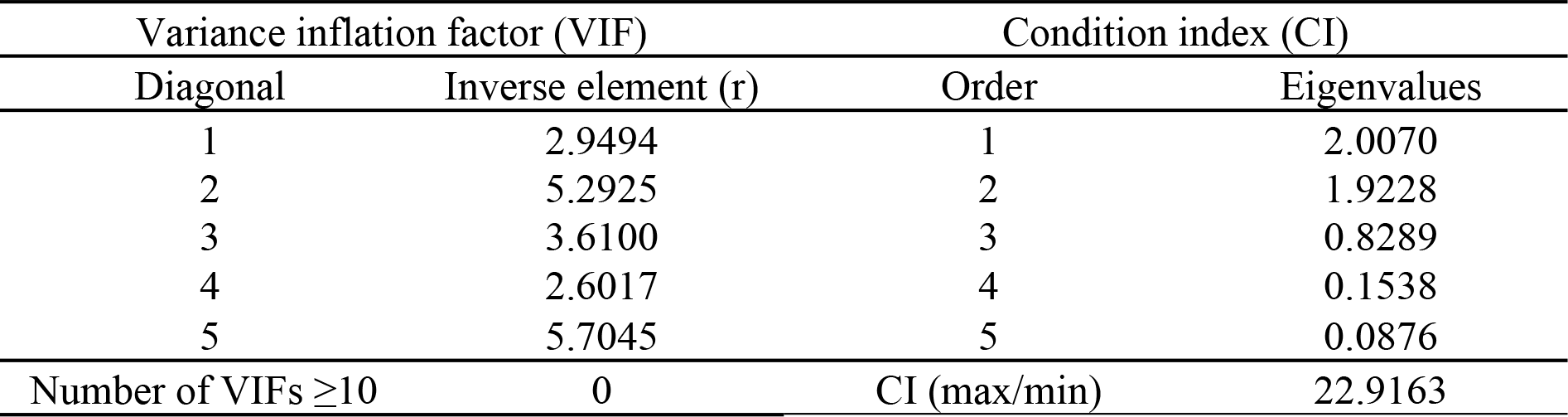
Diagnosis of multicollinearity for traits related to *Spodoptera frugiperda* tolerance in five popcorn varieties (*Zea mays* L.): final larva weight (FW), food intake weight (IW), relative consumption rate (RCR), apparent digestibility (AD) and leaf area consumed (LAC).

| Variance inflation factor (VIF) |  | Condition index (CI) |  |
| --- | --- | --- | --- |
| Diagonal | Inverse element (r) | Order | Eigenvalues |
| 1 | 2.9494 | 1 | 2.0070 |
| 2 | 5.2925 | 2 | 1.9228 |
| 3 | 3.6100 | 3 | 0.8289 |
| 4 | 2.6017 | 4 | 0.1538 |
| 5 | 5.7045 | 5 | 0.0876 |
| Number of VIFs $\geq 10$ | 0 | CI (max/min) | 22.9163 |

Multivariate techniques to select traits related to *S. frugiperda* tolerance in popcorn genotypes were also used by [7, 17] and as in this study, the application of multivariate analysis allowed a significant reduction in the number of traits. This raises the chances of a more effective selection, since the smaller number of traits prevents the effect of interrelationships among them, avoiding redundancy and mistakes in the process of selecting promising genotypes [8].

The selected traits can, at the end of this study, be described as directly related to the damage caused by *S. frugiperda* larvas and can be considered key traits for the tolerance of popcorn genotypes, in that tolerant genotypes will have a smaller leaf area consumed (LAC) and a lower food intake weight (IW) and relative consumption rate (RCR), as well as less apparent digestibility (AD), which will result in a lower final larva weight (FW) [29, 30].

In this study, the use of multivariate techniques for selection and grouping was used as a way to validate both the selection of tolerant genotypes and the influence of the chosen traits on the identification of tolerance.

In multivariate procedures, one of the most widely used techniques is hierarchical clustering [6, 7, 17, 31]. By these methods, genotypes are grouped by a process that is repeated at several levels, establishing a dendrogram, with no predetermined optimal number of groups. For this case, [32] described different forms of representing the clustering structure based on the distance between the genotype pairs, for which UPGMA is the most commonly used method [17, 33, 34].

The relationships between the varieties UEM J1, Zapalote Chico, Composto Márcia, Arachida and Composto Gaúcho can be observed in a graph of the results of the dendrogram based on Mahalanobis’ generalized distance, grouped by the UPGMA method (Figure 1). The high cophenetic correlation coefficient (CCC = 0.93) indicated an optimal fit between the original data and the dissimilarity matrix and high accuracy of the experimental results. The dendrogram shows two main groups, making the discrimination of the tolerance and/or susceptibility of each genotype group possible [1, 5, 23].

**Figure 1.**
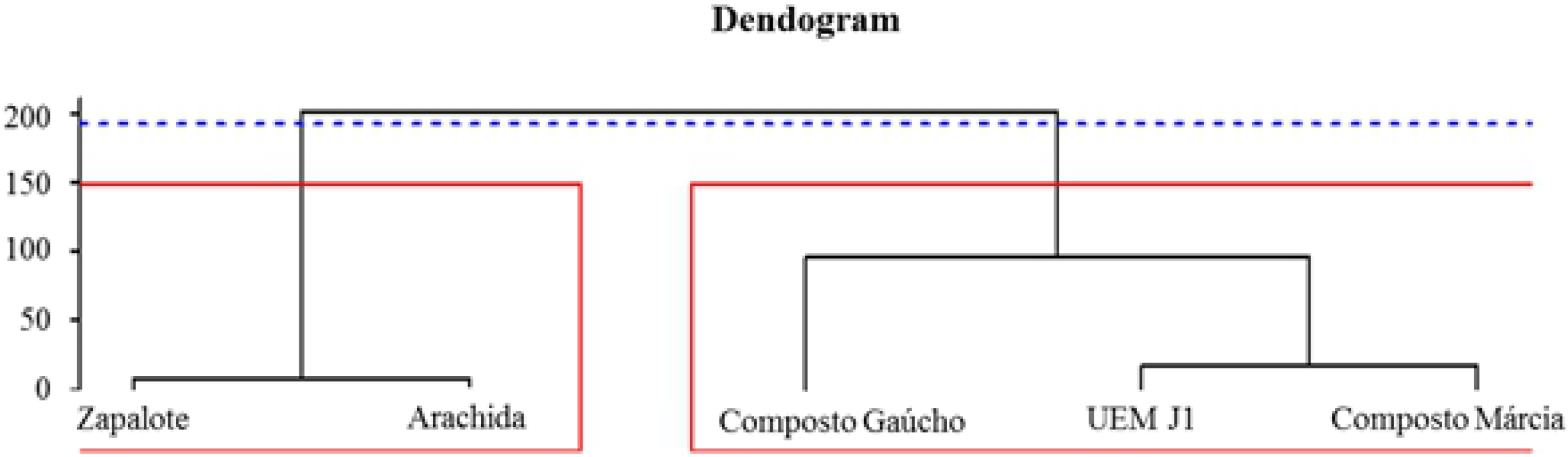
Dendrogram of hierarchical analysis based on Mahalanobis’ generalized distance for the traits grouped by the UPGMA method among five popcorn varieties. Cutline established according to Mojena, (1977).

The first group contained the variety Zapalote Chico, considered by several authors as tolerant to *S. frugiperda* [7, 13, 17], and variety Arachida. The varieties Composto Gaúcho, UEM J1 and Composto Márcia were grouped in the second (Figure 1).

The proximity of the varieties Arachida and Zapalote Chico may suggest tolerance of variety Arachida to *S. frugiperda*, since the grouping was based on the same traits for both varieties. The results show (Table 3) that the varieties Arachida and Zapalote Chico had the lowest values for index of leaf area consumed (LAC), food intake weight (IW) and relative consumption rate (RCR), aside from the low apparent digestibility (AD) and low final larva weight (FW), which indicate tolerance of these two varieties. The cultivar Zapalote Chico is so far the only one treated as tolerant to *S. frugiperda*, so the identification of new tolerant cultivars can be considered a great gain to the breeding since it allows to explore differents sources of tolerance.

**Table 3.** Means of traits related to *Spodoptera frugiperda* tolerance in five popcorn varieties (*Zea mays* L.): final larva weight (FW), food intake weight (IW), relative consumption rate (RCR), apparent digestibility (AD) and leaf area consumed (LAC),

| Genotypes | FW (g) |  | IW (g) |  | RCR (g/g) |  | AD (%) |  | LAC (cm<sup>2</sup>) |  |
| --- | --- | --- | --- | --- | --- | --- | --- | --- | --- | --- |
| UEM J1 | 0.5434 | ab | 2.6850 | bc | 0.0925 | b | 37.3388 | c | 209.6653 | b |
| Zapalote Chico | 0.5306 | b | 2.4318 | d | 0.0780 | d | 39.4875 | bc | 173.8532 | d |
| Composto Márcia | 0.5348 | b | 2.6810 | c | 0.0862 | c | 41.0572 | ab | 209.7829 | b |
| Arachida | 0.5547 | ab | 2.4818 | d | 0.0763 | d | 39.5563 | bc | 186.7935 | c |
| Composto Gaúcho | 0.5678 | a | 2.8366 | a | 0.1088 | a | 43.1369 | a | 222.9237 | a |

Means followed by the same letter in a column do not differ statistically by the Roy and Bose test at 5% probability

Moreover, the performance of variety Composto Gaúcho was the worst, and it was characterized as the most susceptible to *S. frugiperda*, based on the evaluated traits, since the values of FW, IW, RCR, AD and LAC were high (Table 3).

In this study, variety Zapalote Chico was used as check because of its known tolerance of the type antixenosis, i.e., feeding avoidance [13, 35], together with variety Arachida, with antixenosis tolerance as well. This type of tolerance can be inferred from the low food intake rate, low relative consumption and smaller leaf area consumed. In the case of variety Zapalote Chico, the final larva weight was also low, indicating possible antibiosis tolerance.

The dendrogram was confirmed and analyzed in more detail by canonical variable analysis, grouped by Tocher’s clustering (Figure 2), which showed the presence of three distinct groups. Tocher’s optimization is a clustering method based on the formation of groups whose distances within are shorter than the distances between groups. At the end of the process, the number of groups and accessions contained in each group are computed. This method was applied as suggested by [32] and is an important way of determining different groups based on different traits, together with the techniques of dissimilarity analysis and analysis of canonical variables [6].

**Figure 2.**
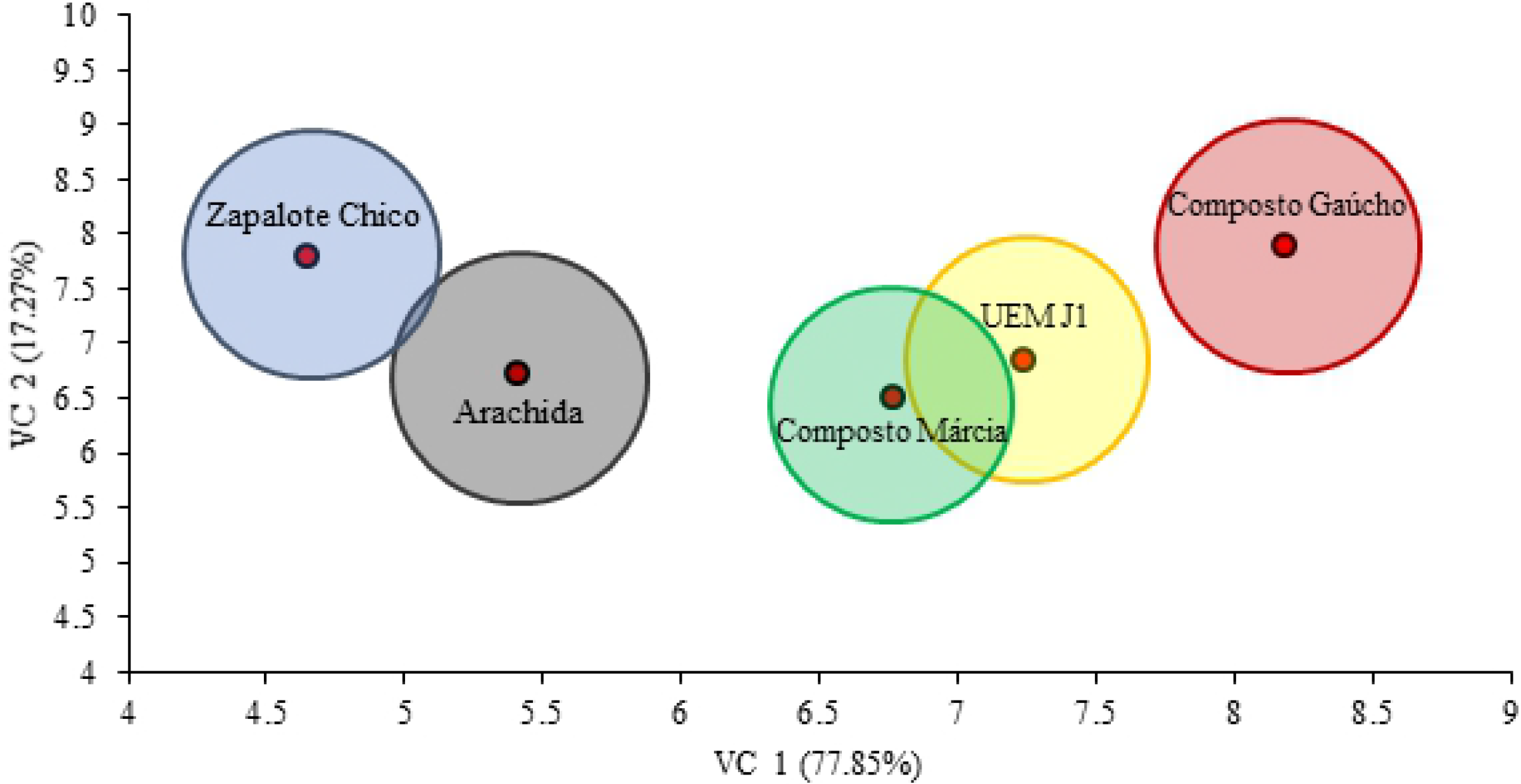
Biplot of canonical variable analysis showing the closest and most distant groups of five popcorn genotypes grouped by Tocher’s clustering.

The analysis of canonical variables explained 95.12% of the total variation between the five traits. When analyzing the dispersion of the scores of the first two canonical variables, there was an agreement with the previous groupings, confirming the results, as well as the choice of the traits used in the study of *S. frugiperda* tolerance. The first canonical variable (VC 1) explained 87.85% of the total variation and the second (VC 2) 25.6% (Figure 2).

Tocher’s grouping method, based on the analysis of canonical variables, grouped the varieties Zapalote Chico and Arachida again. Similarly to the variety Zapalote Chico, considered tolerant by several authors [7, 13, 17] and also in this study, some characteristics of variety Arachida like final larva weight, food intake weight, relative consumption rate, apparent digestibility and leaf area consumed also indicate tolerance. The varieties UEM J1 and Composto Márcia, which, according to the means of the analyzed variables, were moderately tolerant, were contained in the second group. Variety Composto Gaúcho remained isolated from the other evaluated varieties, as the most susceptible to *S. frugiperda*, based on its performance with regard to the analyzed traits (Table 3).

Factor analysis is a multivariate statistical method that has been applied in agronomic studies for a relatively short time [25]. This analysis explains the relationships observed between traits and removes possible redundancies or duplications from a set of correlated phenotypic data [36]. The method allows the selection of relevant traits, exploring the relationships and their variations, apart from generating important information about factors and genotypes [37]. In this study, the factor analysis was directed to the identification of key traits related to tolerance to *S. frugiperda*.

By factor analysis, the traits are replaced by a smaller number of latent traits, called factors. These factors group the traits, so that there is little or no variance within groups but maximum variation between groups [32]. By this technique, associated with the analysis of canonical variables and canonical correspondence, the traits that best discriminate genotypes for a given objective, here *S. frugiperda* tolerance, can be efficiently selected.

In this study, the high commonality (from 0.7012 - A to 0.9912 - RCR), indicated traits with a high relation to the determination of tolerance, confirming the thesis that the selected traits can be considered key traits (Table 4). In the factor analysis, the first factor was determinant for traits IW, FW, and RCR and the second for AD and LAC (Table 4). In this study, estimates above 0.90 were considered for one factor and low estimates for the other factor (Figure 3), which shows a high representativeness of the factor for the respective traits [25]. The estimates of the other traits were intermediate between both determined factors, which despite having a certain degree of contribution to tolerance, were not as expressive as the described traits. In the study of [17], a large number of traits was also reduced, and subsequently divided into two factors by factor analysis.

**Table 4.** Factors and their factorial loads after rotation of the factor axis by the Varimax method for studied traits related to *Spodoptera frugiperda* tolerance in the composite varieties UEM J1, Zapalote Chico, Márcia, Arachida and Gaúcho.

| Variables | Factor score coefficients |  | Loading factors after rotation |  | Commonality |
| --- | --- | --- | --- | --- | --- |
|  | Factor 1 | Factor 2 | Factor 1 | Factor 2 |  |
| LSt | 0.7481 | 0.1367 | 0.6362 | 0.4173 | 0.8784 |
| IW | 0.9849 | 0.1059 | 0.6240 | 0.8694 | 0.9813 |
| FW | 0.9745 | -0.1334 | 0.4302 | 0.9843 | 0.9195 |
| MW | 0.5249 | 0.8217 | 0.1639 | 0.8612 | 0.8508 |
| F | 0.6063 | 0.7794 | -0.1193 | 0.9803 | 0.8751 |
| A | 0.8301 | -0.5578 | 0.9820 | 0.1894 | 0.7012 |
| M | 0.8159 | -0.5627 | 0.9754 | 0.1760 | 0.9823 |
| RCR | 0.9460 | 0.1592 | 0.0645 | 0.9935 | 0.9912 |
| RMR | 0.7935 | 0.4367 | 0.5139 | 0.4968 | 0.7721 |
| RGR | 0.6952 | 0.5956 | 0.0733 | 0.9126 | 0.8381 |
| CEI | -0.8723 | -0.2911 | -0.4136 | -0.8213 | 0.8456 |
| AD | 0.1934 | 0.8863 | 0.8329 | -0.4218 | 0.9716 |
| CED | -0.8346 | 0.4910 | -0.9381 | -0.2400 | 0.9377 |
| LAC | -0.0728 | 0.9870 | -0.6063 | 0.9507 | 0.9625 |

**Figure 3.**
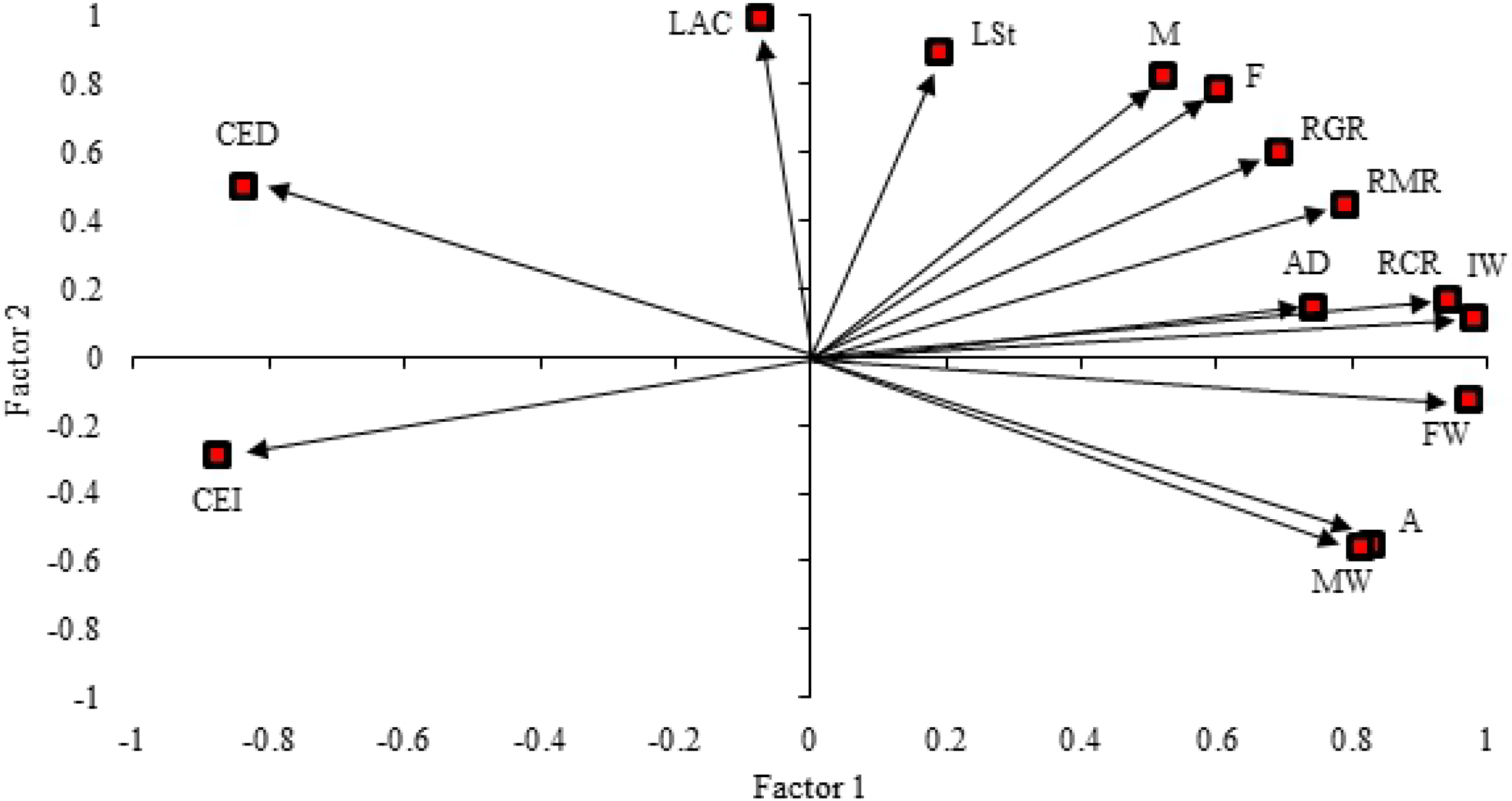
Biplot of factor analysis for traits related to tolerance to *Spodoptera frugiperda* in the composites UEM J1, Zapalote Chico, Márcia, Arachida and Gaúcho. LSt: larval stage duration, IW: food intake weight, FW: final larva weight, MW: mean larva weight, F: feces, A: assimilated food, M: metabolized food, RCR: relative consumption rate, RMR: relative metabolic rate, RGR: relative growth rate, CEI: conversion efficiency of food intake, AD: apparent digestibility, CED: conversion efficiency of the digested food and LAC: leaf area consumed.

According to the factor analysis, the chosen traits are directly correlated with the selection of *S. frugiperda* tolerant genotypes. Thus, we proceeded to the analysis of canonical correspondence.

Canonical correspondence analysis is an exploratory technique to simplify the structure of multivariate data variability, in which the traits are arranged in contingency tables, taking correspondence measures between rows and columns of the data matrix into account. According to [38], correspondence analysis is a method to determine an association system between the elements of two or more sets, to explain the association structure of the factors in question. Thus, graphs were constructed with the principal components of the rows and columns, allowing the visualization of the relationship between the sets, where the proximity of the points referring to the row and the column indicates an association and distance indicates repulsion. According to [39], one of the great advantages of canonical correspondence analysis is that relationships can be detected by this technique that would not have been perceived if the analysis were based on trait pairs. In addition, it is highly flexible in the data traits, since no theoretical model of probability distribution must be adopted. A rectangular matrix containing non-negative data is sufficient, which in the field of breeding, makes it possible to masterfully relate the effects of different traits on specific genotypes.

Canonical correspondence analysis explained 99.67% of the total variation between the genotypes and the respective traits evaluated. The first canonical correspondence axis (CCA 1) explained 99.20% of the total variation and the second axis (CCA 2) accounted for 0.47% (Figure 4). Most of the total variation was already explained in the first CCA, which is desirable, for increasing the accuracy between the cluster and the estimated scores [38, 39].

**Figure 4.**
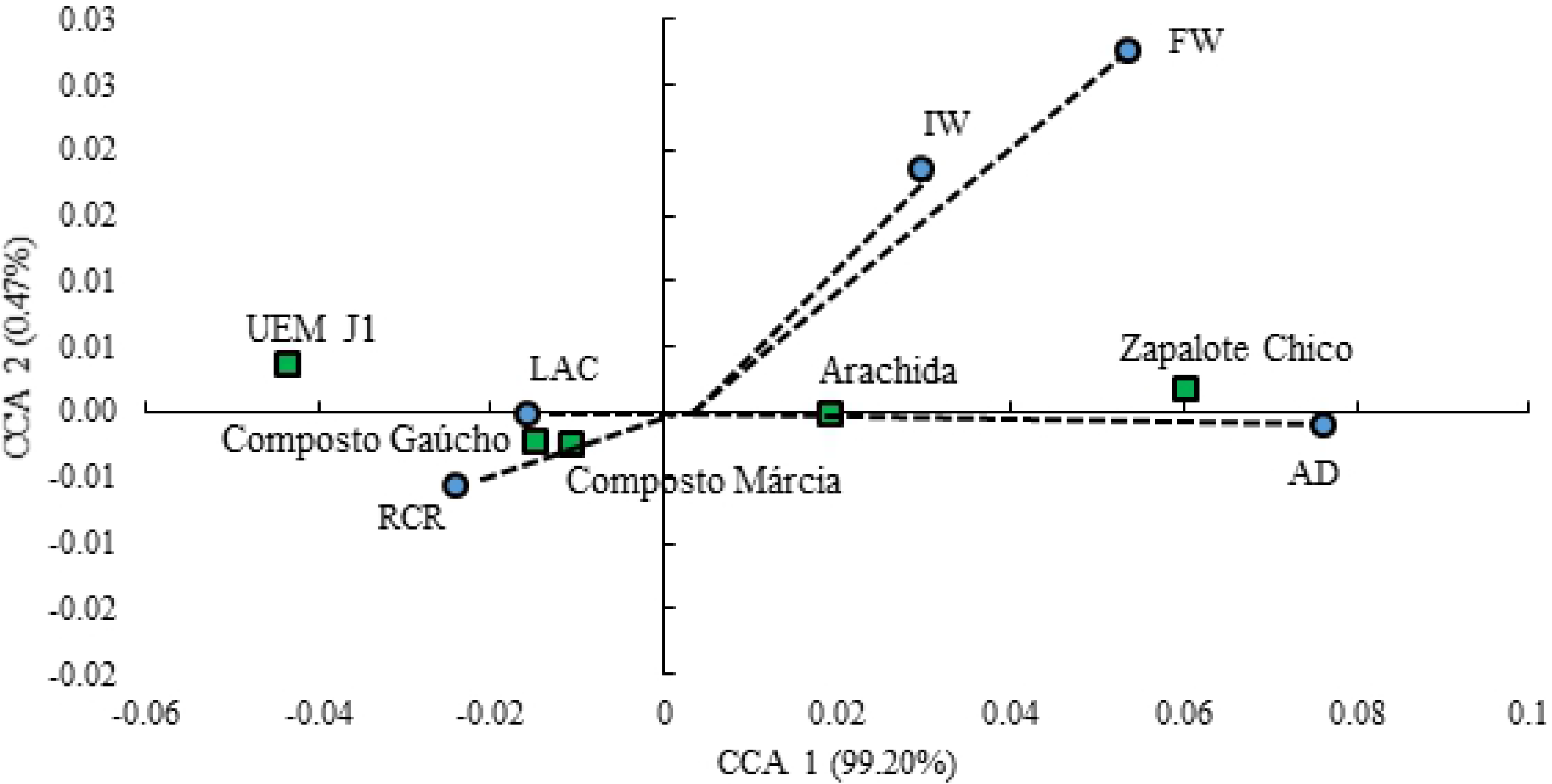
Biplot of canonical correspondence analysis showing the relationship between the five popcorn genotypes and the main explanatory traits of tolerance to *Spodoptera frugiperda*. FW: final larva weight, IW: food intake weight, RCR: relative consumption rate, AD: apparent digestibility, LAC: leaf area consumed.

The biplot of canonical correspondence analysis shows that the traits IW, FW and AD were determinant for the tolerance expressed by the varieties Zapalote Chico and Arachida, in which variable AD had a greater influence, due to its spatial angular proximity in the graph with the said varieties. For the varieties Composto Gaúcho and Composto Márcia, the traits that most contributed to the determination of tolerance or susceptibility were RCR and LAC. For variety UEM J1, the influence of variable LAC was the highest.

In general, the selected traits were efficient in discriminating the genotypes regarding tolerance and susceptibility to *S. frugiperda* by the applied analyses. The identification of key traits in the description of tolerant genotypes will, in future studies, allow greater emphasis on specific traits and consequently a more effective selection regarding tolerance in popcorn genotypes.

## Conclusions

Variety Arachida was identified tolerant to *S. frugiperda* and can be used as a source of favorable alleles for the future development of tolerant popcorn hybrids.

The traits relative consumption rate (RCR), apparent digestibility (AD) and leaf area consumed (LAC) were efficient and considered key traits for the identification of *S. frugiperda* tolerant genotypes.

